# High-Fat Diet-Induced Obesity Impairs Endothelium-Dependent Relaxation via MLCK Activation: Reversal by ML-7 in Rabbits

**DOI:** 10.1101/2025.09.04.674370

**Authors:** Jiao Li, Jiantao Zhou, Qingxv Ha, Junwei Gu, Hui Yuan, Juan Cheng, Xiaojun Zha, Jingjing Teng, Liang Li, Junli Ding

## Abstract

**Objective:** Obesity is an independent risk factor for CVD(cardiovascular diseases) and elevated cardiovascular mortality, though the precise mechanisms remain incompletely elucidated. Impaired endothelium-dependent relaxation is an early manifestation of cardiovascular diseases. This study investigated the impact of HFD(high-fat diet)-induced obesity on endothelium-dependent relaxation function and explored the potential role of MLCK(myosin light chain kinase) signaling.

**Methods:** Forty-five New Zealand White rabbits were randomized into three groups: a control group (normal diet), HFD(high-fat diet) group, and an ML-7 group (HFD plus the MLCK inhibitor ML-7). After 8 weeks of feeding, vascular endothelial function was assessed in vivo by measuring FMD(flow-mediated dilation). Serum lipids and blood glucose levels were determined. Endothelium-dependent vasodilation was evaluated ex vivo using aortic ring assays. The protein expression levels of MLCK and eNOS(endothelial nitric oxide synthase), as well as the phosphorylation level of MLC(myosin light chain), in rabbit arteries were analyzed by Western blot.

**Results:** HFD-fed rabbits developed obesity, dyslipidemia (elevated TC[total cholesterol], LDL-C[low-density lipoprotein cholesterol], and ox-LDL[oxidized low-density lipoprotein]), perirenal adipose tissue deposition, and significant endothelial dysfunction, evidenced by impaired endothelium-dependent relaxation function and morphological endothelial damage. Western blot analysis revealed upregulated MLCK and eNOS expression, along with increased phosphorylation of MLC in the arterial walls of obese rabbits, despite no significant change in NO(nitric oxide) levels compared to controls. Treatment with ML-7 ameliorated endothelium-dependent relaxation function impairment. Furthermore, the inherent limitations of FMD as a non-invasive technique in terms of accuracy and sensitivity for evaluating endothelium-dependent dilation.

**Conclusions:** Obesity-induced endothelial dysfunction arise from visceral fat accumulation, dyslipidemia, oxidative stress, and MLCK overexpression. Notably, MLCK inhibition via ML-7 restores vascular function, highlighting its therapeutic potential in obesity-related vascular injury.

## Introduction

Obesity, as a chronic progressive disease, severely impacts both physical and mental health, reduces quality of life [1], and significantly increases medical costs [2]. It has become a serious challenge for global health systems. Data show that the average BMI across all age groups in China has been continuously rising [3], with the adult obesity prevalence surging from 3.1% in 2004 to 16.4% in 2023 [4]. Asian children, particularly Chinese children, exhibit higher body fat percentages and central obesity characteristics [5–6], which are closely associated with an increased risk of atherosclerosis [7]. Even more alarming is that the 2021 Global Burden of Disease Study revealed China ranks first in the absolute burden of obesity [8]. These statistics highlight the current epidemic status and heavy burden of obesity in China and globally.

Obesity is not only a health risk but also exacerbates inflammation and oxidative stress through various mechanisms, leading to severe complications such as CVD(cardiovascular diseases) [7], metabolic disorders [9], liver disease [10], cancers [11], and joint diseases [12], among which CVD has the most prominent impact. In 2015, 4 million people worldwide died due to high BMI, with over two-thirds of these deaths attributed to CVD [13].AS (atherosclerosis), the pathological basis of most CVD, often begins in childhood through a long and complex process. Childhood obesity and its risk factors can accelerate the progression of AS, leading to early-onset CVD [14–15]. Given the severity and prevalence of AS, it is crucial to study the detection and mechanisms of its initial stage—ED(endothelial dysfunction)[16]. ED includes impaired endothelium-dependent relaxation, compromised barrier function, and vascular inflammation, among others. However, there is currently no universally accepted gold standard for in vivo detection of endothelium-dependent relaxation function. Although ex vivo arterial ring experiments are direct, they are not suitable for clinical use. FMD(flow-mediated dilation) of arteries, as a non-invasive detection method, holds potential for predicting vascular endothelial function [17], but further validation and refinement are still needed.

ED is complex and multifaceted, influenced by a variety of factors including genetics, hypertension, hyperlipidemia, diabetes, obesity, aging, smoking, and physical inactivity [18]. The underlying mechanisms remain to be fully elucidated. Our research has found that upregulation of MLCK(myosin light chain kinase) expression and the subsequent phosphorylation of MLC(myosin light chain) exacerbate ED associated with AS. Notably, ML-7(an MLCK inhibitor) can effectively alleviate this dysfunction [19]. Additionally, ED induced by obesity is closely linked to reduced bioavailability of NO(nitric oxide). NO is a key molecule regulating vasomotor function in endothelial cells, and its production depends on the activation of eNOS(endothelial nitric oxide synthase) [20]. However, when eNOS substrates (L-arginine, oxygen) or cofactors (such as tetrahydrobiopterin) are insufficient, eNOS becomes uncoupled and generates superoxide anions instead [21]. In obese mice, eNOS uncoupling leads to decreased NO production, resulting in aortic vascular dysfunction. The combination of L-arginine and an arginase inhibitor can reverse eNOS uncoupling and fully restore aortic function [22].

This study aims to investigate the changes in vascular endothelium-dependent relaxation function(Acetylcholine-induced aortic ring relaxation rate was used as the primary outcome measure) in HFD(high-fat diet)-induced obese rabbits and their relationship with risk factors such as blood lipids and blood glucose, particularly focusing on the associations with MLCK expression and activity, eNOS expression and activity, and NO levels. These investigations will help us understand the mechanisms underlying obesity-induced endothelium-dependent relaxation dysfunction and validate the reliability of non-invasive FMD as a method for assessing this function.

## Materials and Methods

### 2.1 Animal experimental procedures

After a 2-week acclimatization period, healthy rabbits that met the required age and weight criteria were included in the study. Those not fulfilling the criteria were excluded and replaced in a timely manner to maintain the predetermined sample size in each group. Forty-five male New Zealand White rabbits (purchased from Laifu Animals Breeding Center, Nanjing, China; 3 months old, weighing 2.7 (2.5, 2.8) kg, with an approval number of SCXK(Su)2019-0005, and a certificate number of 202270193) rabbits were randomly allocated into three groups (n=15 per group) using a computer-generated random number table and housed under identical conditions for 8 weeks. HFD Group: Fed a high-fat diet (HFD) containing 0.5% cholesterol, 5% lard, 5% soybean oil, and 89.5% standard chow, and received daily oral gavage of an equal volume of ultrapure water. ML-7 Group: Received the same HFD, in addition to daily oral administration of ML-7 (1 mg/kg; DC Chemicals, China) in ultrapure water. Control Group: Fed a standard chow diet and received daily oral gavage of an equal volume of ultrapure water. The oral gavage procedure was performed as follows: between 9:00 and 11:00 AM daily in a dedicated animal procedure room, the rabbit was restrained, and a silicone catheter was inserted into the esophagus via an oral speculum. After confirming the absence of bubbles at the submerged end of the catheter, the solution was slowly injected using a syringe before the catheter was withdrawn. This procedure was uniformly applied to all groups to isolate the effects of the drug from those of gavage-related stress. After the 8-week experimental period, all animals were euthanized for subsequent tissue collection and analysis. The experimental protocol was approved by the Ethics Committee of Anhui Medical University (Hefei, China).

### 2.2 Tissue Harvesting and Processing

After euthanasia, the thoracic aorta was rapidly isolated through thoracotomy and placed in 4°C Krebs solution (maintained on ice), continuously aerated with a gas mixture of 95% O₂ and 5% CO₂. Adipose and connective tissues surrounding the aorta were carefully removed using ophthalmic instruments. A 6–8 mm segment of the aortic ring was excised for ex vivo vascular ring experiments, while the remaining tissue was preserved for subsequent analyses, including ORO(Oil Red O) staining and paraffin sectioning. PRAT(perirenal adipose tissue), situated between the renal capsule and renal fascia, was collected and weighed[23].

### 2.3 Lipid Profile and Blood Glucose Measurements

Serum was collected from the auricular veins of New Zealand white rabbits. After clotting for 2 h at room temperature, samples were centrifuged (3500 × g, 15 min, 4°C) to isolate serum. The supernatant was aliquoted into sterile tubes, wrapped in aluminum foil to protect from light, and stored at -80°C to avoid freeze-thaw degradation. Serum glucose(Glu) and lipid profiles — including TC(total cholesterol), TG(triglycerides), LDL-C(low-density lipoprotein cholesterol), HDL-C(high-density lipoprotein cholesterol), ox-LDL(oxidized low-density lipoprotein), and VLDL-C(very low-density lipoprotein cholesterol) — were quantified using commercial ELISA kits (Shanghai Jianglai Biological Technology Co., Ltd and Jiangsu Jingmei Biological Technology Co., Ltd) following manufacturer protocols, with internal controls to ensure reliability.

### 2.4 ORO staining

The aortic tissue was dissected and stained with ORO solution (prepared by dissolving 0.25 g ORO in 50 mL of 60% isopropanol and diluted with distilled water at a 3:2 ratio before staining) for 20 minutes. After staining, the tissue was washed with 60% isopropanol to remove unbound dye. The staining results were qualitatively analyzed by imaging to assess the distribution of fat infiltration in the aorta.

### 2.5 H&E(Hematoxylin and Eosin) Staining

The thoracic aortic specimens were trimmed into 5 mm thick sections, fixed overnight in 4% formaldehyde, dehydrated through an ethanol gradient, and embedded in paraffin. The sections were subjected to H&E staining and observed under an optical microscope (Leica, Wetzlar, Germany) to examine the arterial wall structure.

### 2.6 WB(Western Blotting)

The aortic tissue was washed three times with TBS(tris-buffered saline) and lysed using RIPA (pre-cooled Radioimmunoprecipitation assay) buffer, followed by centrifugation at 18,800 g for 30 minutes at 4 ° C. Total protein concentration was determined using a Micro-bicinchoninic acid kit (Pierce, Rockford, IL, USA), and a standard curve was plotted. Equal amounts of protein samples were separated by 10% SDS-PAGE(sodium dodecyl sulfate – polyacrylamide gel electrophoresis) and transferred to PVDF(polyvinylidene fluoride) membranes, which were then blocked with 5% non-fat TBST(tris-buffered saline with Tween-20) (containing 0.1% Tween-20) at room temperature for 2 hours. The membranes were incubated with the following primary antibodies: MLCK (1:1000, Sigma, M7905), p-MLC(phosphorylated myosin light chain)(S20) (1:5000, Abcam, ab2480), MLC (1:1000, Santa Cruz, sc-365243/sc-293136), eNOS (1:1000, Immunoway, YT3174), and β-actin (1:10000, Santa Cruz, sc-47778).

### 2.7 Measurement of NO in Rabbit Arterial Walls

The treated arterial tissue was homogenized and centrifuged, and the supernatant was collected and kept on ice for analysis. The total NO concentration in the arterial wall was measured using a commercial assay kit (Elabscience, E-BC-K035-M).

### 2.8 Non-invasive Ultrasound Assessment of Endothelial Function in the Left External Iliac Artery

After shaving the abdominal and left groin areas of rabbits, FMD of the left external iliac artery was assessed using transcutaneous ultrasound. Rabbits were anesthetized with 3% sodium pentobarbital solution (30 mg/kg) via the marginal ear vein, placed in a supine position, and ensured airway patency. Ultrasound gel was applied to the left groin. A GE Vivid E9 (USA) ultrasound system with a 9L8 MHz probe was used to locate the bifurcation of the left common iliac artery and track the external iliac artery, obtaining longitudinal images with the beam perpendicular to the arterial wall. The baseline end-diastolic diameter (D0) of the external iliac artery was measured. A cuff was then inflated to 240 mmHg for 5 minutes at the left hindlimb, and the maximum end-diastolic diameter (D1) was measured after sudden deflation. The procedure was repeated three times, and the average value was calculated. FMD (%) was determined as (D1-D0)/D0 × 100% to evaluate endothelium-dependent dilation [19].

### 2.9 Analysis of Ex Vivo Arterial Ring Relaxation Function

Thoracic aortic rings from rabbits were preserved with intact endothelium. Briefly, arterial rings were suspended in a 37°C Krebs solution bath (continuously aerated with 95% O₂ and 5% CO₂), and tension was recorded using a BL-420 F system (Chengdu Taimeng Science and Technology Co., Ltd., China). After equilibration at 2 g resting tension for 45 minutes, rings were pre-activated with 80 mM potassium chloride to assess functional integrity and enhance contractile response, followed by pre-contraction with 1 μmol/L phenylephrine. Upon reaching stable contraction, cumulative concentrations of acetylcholine (ACh) or sodium nitroprusside (SNP) (10^-3^ to 10 mol/L) were added. Relaxation was expressed as a percentage of the pre-contraction amplitude.

### 2.10 Analysis

Data were analyzed using SPSS 20.0 and graphed with GraphPad Prism. WB band intensities were quantified in Fiji (ImageJ) and expressed as target protein/internal control ratios. Categorical data are presented as n (%) and compared by Chi-square test. All continuous data were tested for normality and homogeneity of variance. Normally distributed data are expressed as mean ± SD and analyzed by one-way ANOVA. If the overall difference was significant (P < 0.05), LSD or Tamhane’s T2 post hoc test was applied depending on whether variances were equal or not. Non-normally distributed data are expressed as median (P25, P75) and analyzed by Kruskal–Wallis test, followed by Dunn’s post hoc test if significant. Bivariate correlation analysis assessed the correlation of risk factors (e.g.,PRAT, blood lipids) with endothelium-dependent vasodilation. A p-value < 0.05 was considered statistically significant.

### 2.11 Blinding

The randomizer knew groups but did not perform experiments. The investigator administering ML-7 was not involved in data collection. All outcome assessments and data analyses were performed by investigators blinded to the group allocation.

## Results

### 3.1 Successful Establishment of High-Fat Diet-Induced Obesity Model

Initial body weights showed no significant differences among groups. From week 2 onward, both the HFD and ML-7 groups exhibited significantly higher body weights compared to the control group (P < 0.05), with no difference between the two groups. This trend persisted until the end of the experiment. By week 8, the HFD group showed a 27.85% increase in body weight compared to the control group (>25%), meeting the criteria for an obesity model[24].

### 3.2 PRAT Deposition and Dyslipidemia in Obese Rabbits with Limited Efficacy of ML-7 Intervention

Compared to the control group, both HFD and ML-7 groups showed significant increases in perirenal fat, with no significant difference between the two groups (Figure 1A). After 8 weeks of feeding, TC, LDL-C and ox-LDL levels in the HFD and ML-7 groups were significantly higher than those in the control group, with ML-7 group ox-LDL levels exceeding those of the HFD group (P < 0.05). Glucose levels were significantly elevated in the HFD group relative to controls, but ML-7 administration significantly lowered them (Table 1). The results indicate that HFlD-fed rabbits developed obesity with pronounced visceral fat deposition, dyslipidemia (increased TC, LDL-C, and ox-LDL), and hyperglycemia, contributing to a higher risk of atherosclerosis and cardiovascular disorders. However, although ML-7 alleviated hyperglycemia, it did not improve the remaining metabolic parameters.

**Figure 1.**
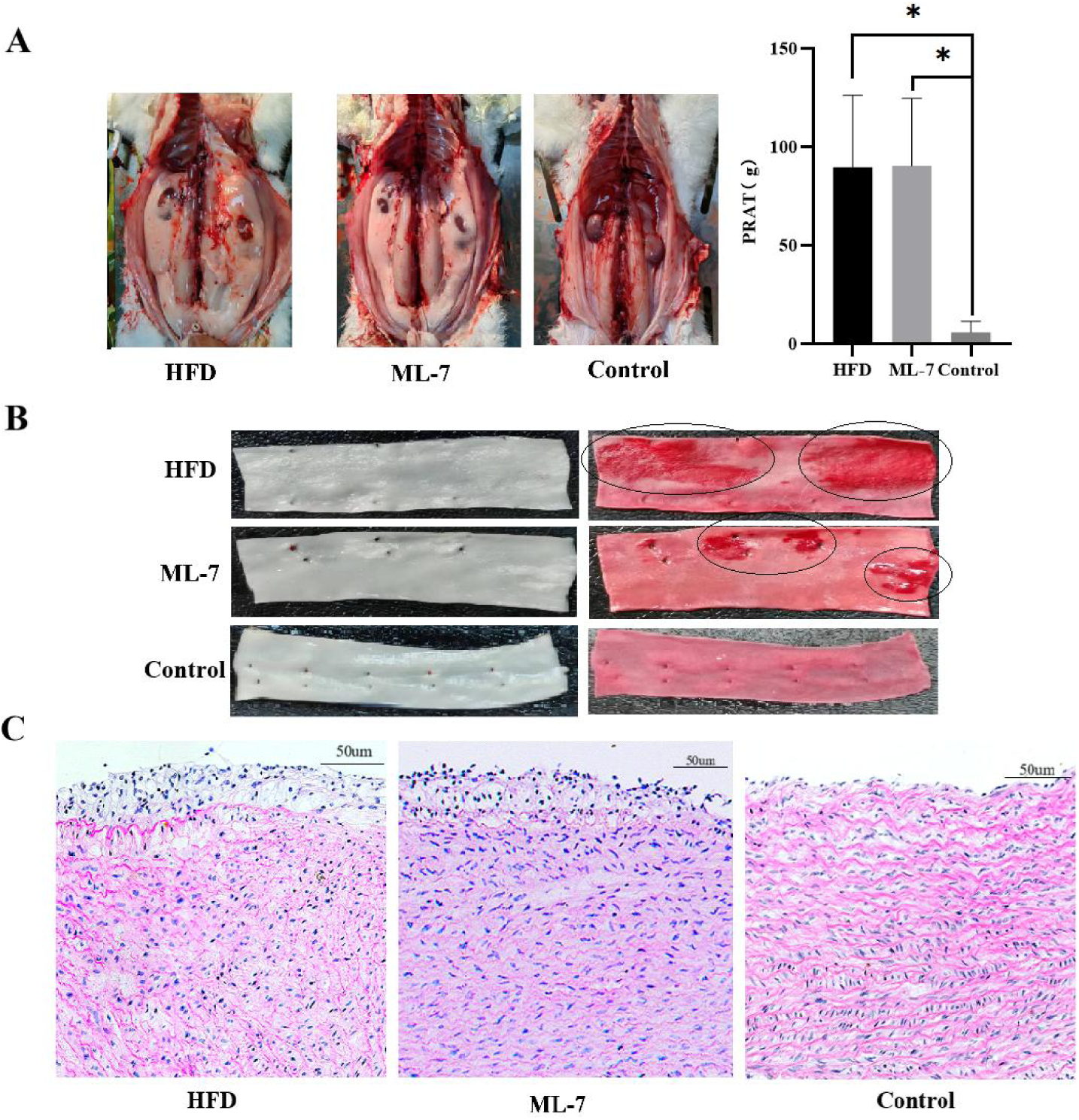
Comparison of PRAT(perirenal adipose tissue)(n=15, N=45)(A), arterial wall appearance, Oil Red O staining(n=9, N=27)(B) and Hematoxylin and Eosin staining(n=3, N=9)(C) of rabbits in different groups. *P<0.05, VS Control.

**Table 1.**
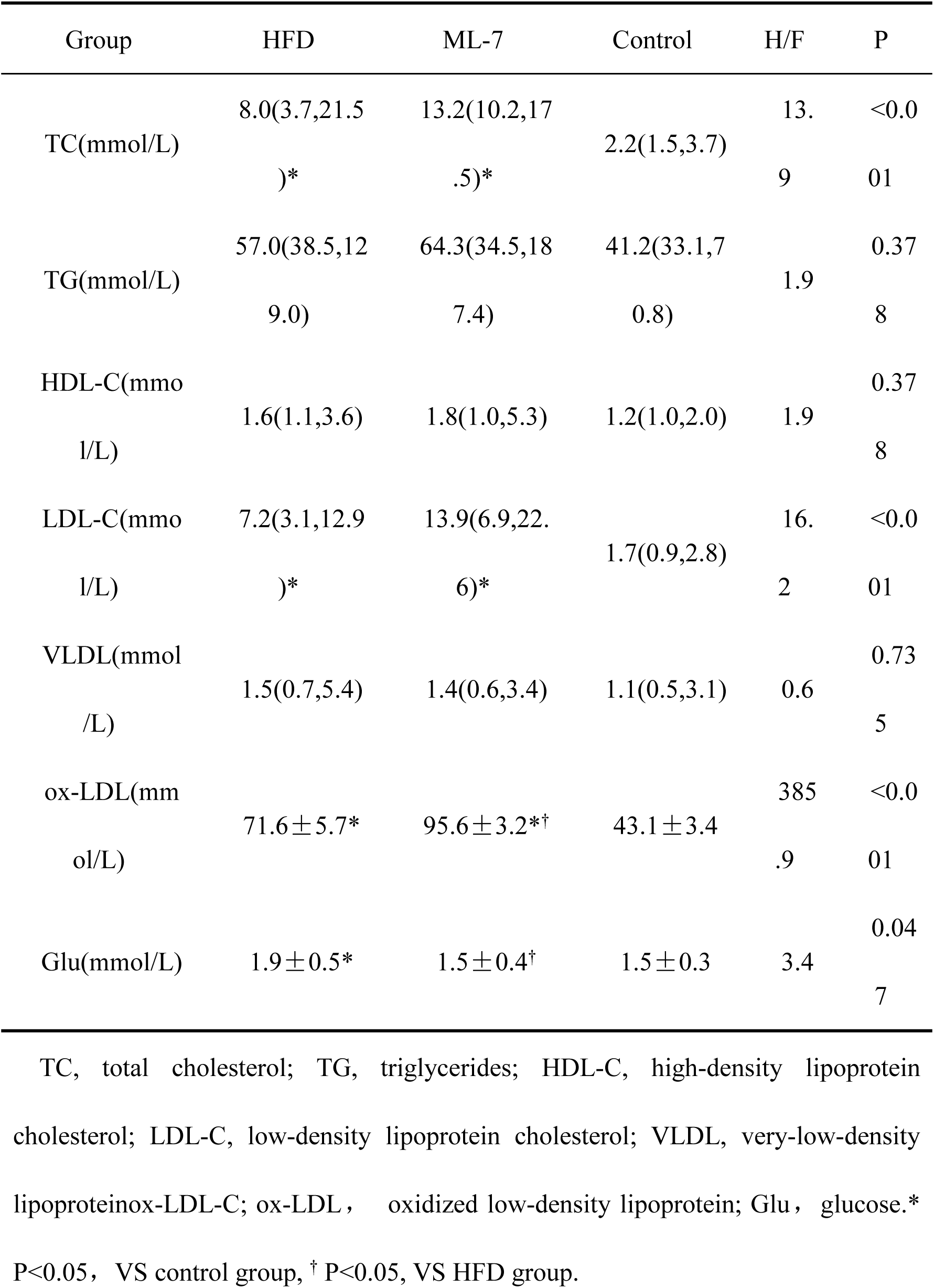
Comparison of blood lipid and glucose levels of experimental rabbits in each group(n=10, N=30).

| Group | HFD | ML-7 | Control | H/F | P |
| --- | --- | --- | --- | --- | --- |
| TC(mmol/L) | 8.0(3.7,21.5)<br>)* | 13.2(10.2,17<br>.5)* | 2.2(1.5,3.7) | 13.<br>9 | <0.0<br>01 |
| TG(mmol/L) | 57.0(38.5,12<br>9.0) | 64.3(34.5,18<br>7.4) | 41.2(33.1,7<br>0.8) | 1.9 | 0.37<br>8 |
| HDL-C(mmo<br>l/L) | 1.6(1.1,3.6) | 1.8(1.0,5.3) | 1.2(1.0,2.0) | 1.9 | 0.37<br>8 |
| LDL-C(mmo<br>l/L) | 7.2(3.1,12.9<br>)* | 13.9(6.9,22.<br>6)* | 1.7(0.9,2.8) | 16.<br>2 | <0.0<br>01 |
| VLDL(mmol<br>/L) | 1.5(0.7,5.4) | 1.4(0.6,3.4) | 1.1(0.5,3.1) | 0.6 | 0.73<br>5 |
| ox-LDL(mm<br>ol/L) | 71.6±5.7* | 95.6±3.2*<sup>†</sup> | 43.1±3.4 | 385<br>.9 | <0.0<br>01 |
| Glu(mmol/L) | 1.9±0.5* | 1.5±0.4<sup>†</sup> | 1.5±0.3 | 3.4 | 0.04<br>7 |

TC, total cholesterol; TG, triglycerides; HDL-C, high-density lipoprotein cholesterol; LDL-C, low-density lipoprotein cholesterol; VLDL, very-low-density lipoprotein; ox-LDL-C; ox-LDL, oxidized low-density lipoprotein; Glu, glucose.\* P<0.05, VS control group, <sup>†</sup> P<0.05, VS HFD group.

### 3.3 Results of aortic ORO staining and HE staining in each group

At 8 weeks, visible differences in aortic wall lesions were observed among groups. Compared to the control group, both the HFD and ML-7 groups exhibited significant ORO staining for plaques, although ML-7 intervention reduced plaque number and area (Figure 1B). HE staining revealed significant structural damage in the aortic endothelium of obese rabbits, exhibiting marked thickening of the aortic intima accompanied by abundant foam cells, lipid deposition, and disorganized smooth muscle cells in the medial layer. Overall, these pathological changes were partially alleviated in the ML-7 group. (Figure 1C).

### 3.4 In Vivo Iliac Artery Ultrasound Reveals Impaired Endothelium-Dependent Dilation in Obese Rabbits with Limited Efficacy of ML-7 Intervention

FMD of the external iliac artery, assessed by transcutaneous ultrasound, showed that both the HFD group and the ML-7 group exhibited significantly lower endothelium-dependent dilation compared to the control group (P < 0.05). However, there was no significant difference in FMD between the ML-7 group and the HFD group (P > 0.05)(Figure 2).

**Fig. 2.**
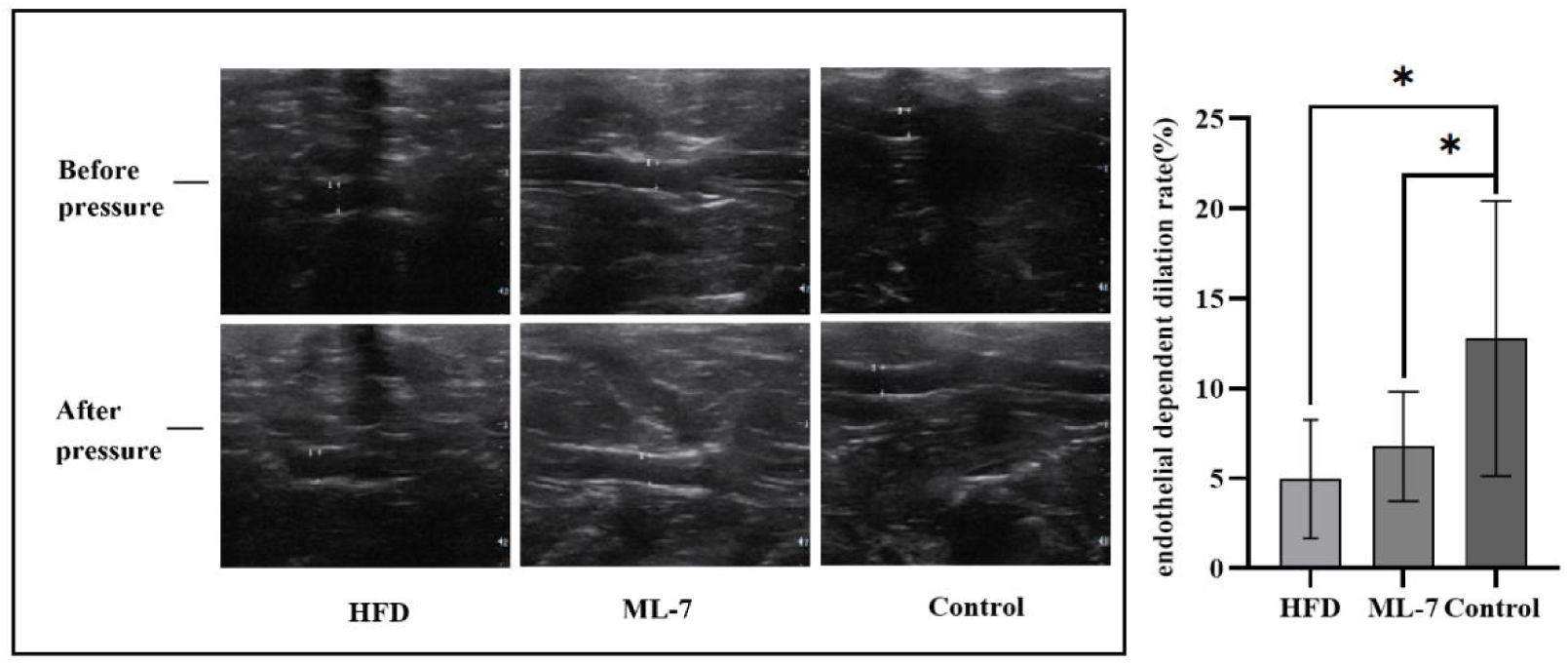
Comparison of endothelial dependent dilation rate of external iliac artery in rabbits fed with different groups(n=8, N=24). Endothelial dependent dilation rate (%) = (D1 − D0) / D0 × 100, where D0 is baseline diameter (before pressure) and D1 is after-pressure diameter of the left external iliac artery. *P<0.05, VS Control.

### 3.5 ML-7 Ameliorates Endothelium-Dependent Relaxation Dysfunction in HFD-Induced Obese Rabbits

HFD group showed reduced relaxation response to 0.1, 1, and 10 μM Ach compared to controls (P < 0.05). ML-7 treatment improved relaxation in high-fat diet-induced obese rabbits at 1 and 10 μM Ach (P < 0.05), but had no significant effect at other concentrations (P > 0.05)(Figure 3A). No significant differences were observed among the three groups at all SNP concentrations (P > 0.05) (Figure 3B). These results indicate that the impaired relaxation function in HFD-induced obese rabbits is endothelium-dependent. ML-7 partially ameliorates this dysfunction.

**Fig. 3.**
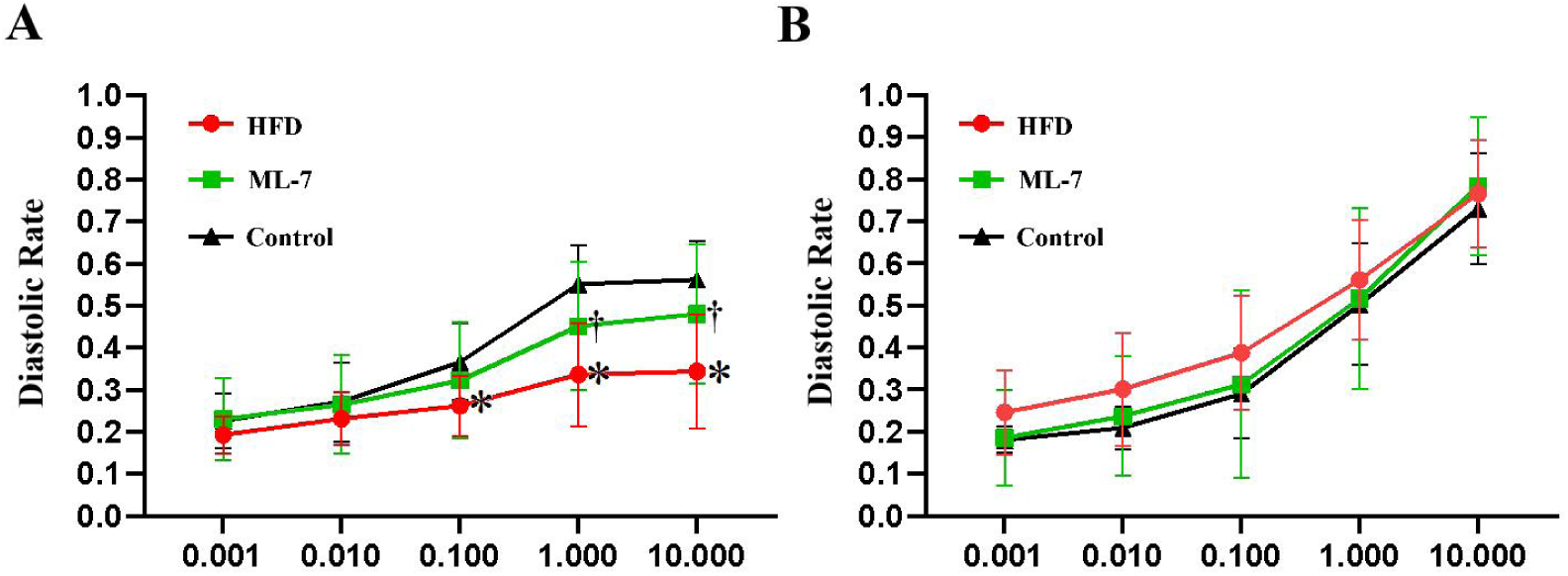
Relaxation response to ACh (A) and SNP (B) in the thoracic aortic rings from rabbits in the three groups(n=7, N=21). Aortic rings were exposed to cumulative doses of ACh and SNP. Values represent percentage of relaxation relative to phenylephrine precontraction (1 mmol/L). ACh, acetylcholine; SNP, sodium nitroprusside. * P< 0.05, VScontrol group, ^†^ P< 0.05, ML-7 group vs HFD group.

### 3.6 Correlation Analysis of Ex Vivo Arterial Ring Relaxation Rates with Other Indicators

Correlation analysis revealed that perirenal fat was significantly positively correlated with body weight and negatively correlated with relaxation rates at Ach 0.1, Ach 1, and Ach 10 (P < 0.05). TC, LDL-C and ox-LDL levels showed a positive correlation with body weight and perirenal fat (Table 2). No significant correlations were observed between body weight, PRAT, blood lipid levels, and relaxation rates at all SNP concentrations (P > 0.05).

**Table 2.** Correlation analysis of diastolic rate of isolated arterial rings with blood lipids, glucose, body weight and PRAT.

|  |  |  |  | Diastolic rate |  |  |  |  |
| --- | --- | --- | --- | --- | --- | --- | --- | --- |
|  |  | weight | PRAT | Ach0.001 | Ach0.01 | Ach0.1 | Ach1 | Ach10 |
| weight | r | 1 | .812* | -0.136 | -0.203 | -0.326 | -0.34 | -0.365 |
|  | P | - | <0.001 | 0.556 | 0.376 | 0.149 | 0.132 | 0.104 |
| PRAT | r | .812* | 1 | -0.412 | -0.416 | -.611* | -.710* | -.727* |
|  | P | <0.001 | - | 0.063 | 0.061 | 0.003 | <0.001 | <0.001 |
| Glu | r | 0.287 | 0.164 | -0.293 | -0.291 | -0.31 | -0.304 | -0.317 |
|  | P | 0.125 | 0.386 | 0.223 | 0.227 | 0.197 | 0.205 | 0.186 |
| TC | r | .387* | .372* | -0.085 | -0.091 | -0.182 | -0.227 | -0.148 |
|  | P | 0.038 | 0.047 | 0.738 | 0.718 | 0.47 | 0.365 | 0.558 |
| TG | r | 0.267 | 0.271 | 0.041 | -0.051 | -0.126 | -0.215 | -0.181 |
|  | P | 0.154 | 0.148 | 0.867 | 0.835 | 0.608 | 0.378 | 0.458 |
| HDL-C | r | 0.267 | 0.271 | 0.041 | -0.051 | -0.126 | -0.214 | -0.181 |
|  | P | 0.154 | 0.148 | 0.867 | 0.835 | 0.608 | 0.378 | 0.459 |
| LDL-C | r | .470* | .429* | 0.066 | 0.009 | -0.084 | -0.235 | -0.213 |
|  | P | 0.009 | 0.018 | 0.789 | 0.971 | 0.731 | 0.333 | 0.382 |
| ox-LDL | r | .709* | .639* | 0.049 | -0.009 | -0.182 | -0.323 | -0.263 |
|  | P | <0.001 | <0.001 | 0.842 | 0.97 | 0.456 | 0.177 | 0.277 |
| VLDL | r | -0.045 | 0.098 | -0.203 | -0.099 | -0.161 | -0.297 | -0.336 |
|  | P | 0.816 | 0.612 | 0.419 | 0.695 | 0.523 | 0.232 | 0.173 |

PRAT, perirenal adipose tissue; TC, total cholesterol; TG, triglycerides; HDL-C, high-density lipoprotein cholesterol; LDL-C, low-density lipoprotein cholesterol; VLDL, very-low-density lipoprotein; ox-LDL-C; ox-LDL, oxidized low-density lipoprotein; Glu, glucose.\*P<0.05.

### 3.7 HFD-induced obese rabbits exhibited significantly increased arterial wall MLCK expression and MLC phosphorylation levels, which were reversed by ML-7 intervention

Western blot analysis showed that the HFD group had significantly higher aortic MLCK expression and p-MLC/MLC levels than the control and ML-7 groups (P < 0.05), with no significant differences among other groups (P > 0.05)(Figure 4A).

**Fig. 4.**
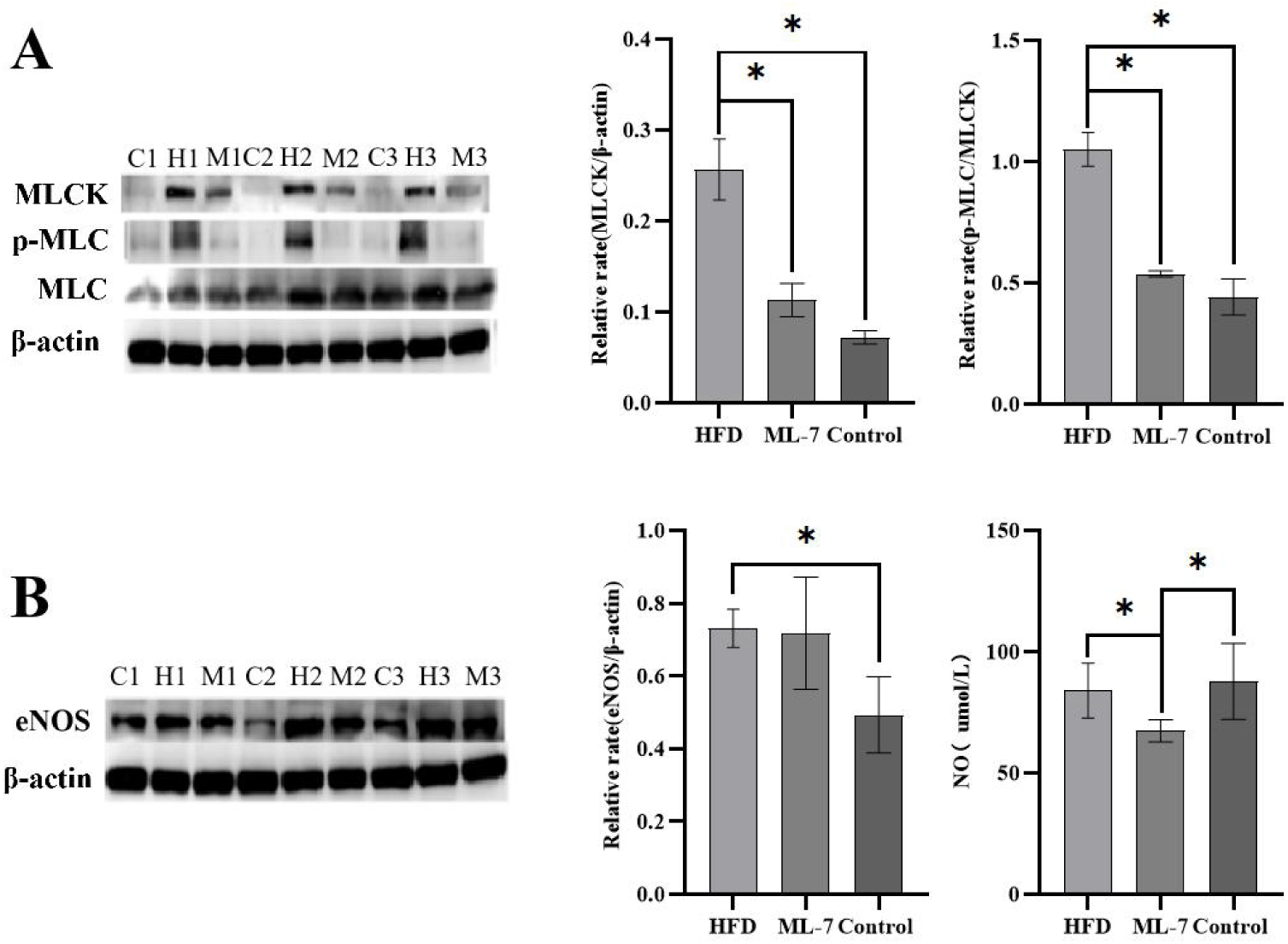
Expression of arterial MLCK, eNOS, NO and phosphorylation levels of MLC in each group(n=3, N=9, with three independent replicates). * P<0.05. C, control group; H, HFD group; M, ML-7 group; MLCK, myosin light chain kinase; MLC, myosin light chain; p-MLC, phosphorylated myosin light chain; eNOS, endothelial nitric oxide synthase.

### 3.8 High-Fat Diet-Induced Obesity Increases Aortic eNOS Expression in Rabbits, but NO Levels Show No Significant Intergroup Differences

Western blot analysis revealed that at 8 weeks, the HFD group exhibited significantly higher aortic eNOS expression compared to the control group (P < 0.05), but NO levels showed no significant difference, potentially due to eNOS uncoupling caused by oxidative stress. The ML-7 group had significantly lower NO levels than the control and HFD groups, possibly due to higher oxidative stress levels (Figure 4B).

## Discussion

Obesity can independently promote the development of CVD and increase CVD-related mortality, distinct from other risk factors [7]. While the precise pathophysiology remains incompletely elucidated, our study investigated obesity-induced vascular dysfunction by focusing on its earliest manifestation — ED. Our findings demonstrate that obesity triggers structural and functional vascular impairment in rabbits, characterized by arterial endothelial damage and compromised endothelium-dependent vasorelaxation. Mechanistically, this obesity-associated ED appears to be driven by a multifactorial interplay of dyslipidemia, oxidative stress, visceral adiposity, and upregulation of the MLCK pathway with consequent elevation in MLC phosphorylation levels. Notably, the amelioration of ED following ML-7 treatment not only validates the pathogenic role of MLCK in obesity-related vascular injury but also provides a theoretical foundation for the early detection and intervention of obesity-related vascular dysfunction in clinical practice.

LDL and its oxidized form (ox-LDL) are well-established pathogenic drivers of atherosclerosis, where LDL-C serves as the classical biomarker of LDL[25]. Consistent with this paradigm, our study demonstrated that obese rabbits exhibited significantly elevated levels of both LDL-C and ox-LDL. The excessive retention of ox-LDL in the subendothelial space and its subsequent uptake by macrophages lead to foam cell formation, promoting ED. Concurrently, ED facilitates the transport of LDL to the subendothelial space, where it is further oxidized into ox-LDL, creating a vicious cycle. Notably, the marked rise in ox-LDL levels observed in our model not only reflects enhanced LDL oxidation but also implies a state of systemic oxidative stress. Multiple enzymatic pathways contribute to this process, including vascular nicotinamide adenine dinucleotide phosphate oxidase, myeloperoxidase activity, mitochondrial dysfunction, and uncoupled eNOS — all of which generate reactive oxygen species and reactive nitrogen species that oxidize LDL [26–27]. Importantly, high-fat diets not only induce hyperlipidemia but also enhance oxidative stress and inflammation, perpetuating a vicious cycle [28], which aligns with our findings.

Visceral and ectopic fat are key drivers of adverse cardiometabolic outcomes in obese patients, with visceral fat closely linked to arterial function [29]. PRAT, a major component of visceral fat, secretes adipokines and pro-inflammatory cytokines, and its thickness positively correlates with visceral fat area [30–31]. Recent studies suggest that PRAT plays a significant role in maintaining cardiovascular and renal homeostasis[23]. Our experimental findings demonstrate that diet-induced obese rabbits developed significant PRAT accumulation. Importantly, PRAT correlated positively with weight but negatively with aortic relaxation at all tested Ach concentrations (0.1, 1, and 10 uM; P < 0.05). Furthermore, serum lipid parameters - including TC, LDL-C, and ox-LDL - exhibited positive correlations with both weight and PRAT volume. These findings collectively suggest a pathogenic interplay between adiposity, dyslipidemia, and vascular dysfunction, with PRAT accumulation showing particularly strong associations with impaired endothelium-dependent vasorelaxation. Emerging evidence suggests PRAT may influence cardiovascular function through multiple pathways, including: (1) neural reflex modulation, (2) adipokine secretion, and (3) fat-kidney crosstalk[30]. Mechanistically, PRAT expansion may promote endothelial dysfunction via increased tetrahydrobiopterin oxidation and subsequent superoxide overproduction, ultimately reducing NO bioavailability [32]. However, the exact pathogenic mechanisms require further elucidation.

This study demonstrates that ED in obese rabbits is associated with MLCK overexpression, increased MLC phosphorylation, dyslipidemia, and perirenal fat deposition, while ML-7 intervention improves endothelium-dependent relaxation. Notably, ML-7-treated rabbits demonstrated paradoxical increases in ox-LDL (P<0.05 vs control/HFD) accompanied by decreased NO content, indicating a potential ML-7-induced oxidative stress response. This phenomenon may be mediated through eNOS uncoupling, as oxidative stress is known to disrupt the eNOS dimerization state and shift its function from NO production to superoxide generation [33].

Additionally, FMD measurements in the left external iliac artery showed only partial agreement with the ex vivo aortic ring experiments (the gold standard for assessing endothelial dilation). This discrepancy may arise from two factors: first, the inherent limitations of FMD as a non-invasive technique in terms of accuracy and sensitivity for evaluating endothelium-dependent dilation; second, the anatomical differences between measurement sites (external iliac artery vs thoracic aorta) that may affect result comparability. Although atherosclerosis is a systemic disease, significant heterogeneity exists in plaque characteristics and functional alterations across different vascular beds [34].

In summary, obesity leads to arterial endothelial cell damage and impaired endothelium-dependent relaxation in rabbits. Obesity-related ED is closely associated with dyslipidemia, oxidative stress, visceral fat deposition, MLCK overexpression, and increased MLC phosphorylation. ML-7 intervention improves endothelium-dependent relaxation in obese rabbits, providing a theoretical foundation for the early detection and intervention of obesity-related vascular dysfunction in clinical practice.

## Acknowledgments

We thank Qingxv Ha, Jiantao Zhou, Junli Ding from the research study team for their excellent administrative support.

## Supplemental Material

Tables S1

Major Resources Table

## Highlights

Highlight 1: Establishes a Link Between Metabolic Phenotype and Vascular Dysfunction

This study delineates a clear path from HFD-consumption to endothelial injury, demonstrating that the development of obesity, dyslipidemia (elevated TC, LDL-C, and ox-LDL), and significant perirenal adipose tissue deposition are directly associated with the impairment of endothelium-dependent relaxation, a key early event in cardiovascular disease.

Highlight 2: Identifies MLCK as a Novel Mechanistic Hub and Therapeutic Target

The research provides crucial evidence that vascular MLCK overexpression and subsequent MLC phosphorylation are key drivers of obesity-induced endothelial dysfunction, a pathology that occurs despite concurrent dyslipidemia and visceral adiposity. Notably, pharmacological inhibition of MLCK with ML-7 effectively restored vascular function, highlighting its therapeutic potential.

## Non-standard Abbreviations and Acronyms

MLCK: myosin light chain kinase
eNOS: endothelial nitric oxide synthase
MLC: myosin light chain
p-MLC: phosphorylated myosin light chain
FMD: flow-mediated dilation
ED: endothelial dysfunction
Ach: acetylcholine
SNP: sodium nitroprusside
PRAT: perirenal adipose tissue

